# A Differentiable dFBA Simulator for Scalable Bayesian Inference over Microbial Metabolic Models

**DOI:** 10.64898/2026.05.05.722888

**Authors:** Tomek Diederen, Charlotte Merzbacher, Martin Patz

## Abstract

Medium optimisation for bioprocess design remains challenging and costly: fermentation recipes typically contain ten or more components, the design space expands combinatorially as ingredients are added, and each batch experiment requires over 24 hours. High-throughput 96-well plate screening can reduce experimental cost, but extracting actionable predictions from growth curves requires a mechanistic model that links medium composition to cellular metabolism. In this paper, we present a differentiable simulator for dynamic flux balance analysis (dFBA) that enables scalable Bayesian inference over microbial metabolic models. A distinguishing feature is that inference is driven entirely by OD600 measurements, a simple optical proxy for biomass, without substrate or product assays; internal fluxes, substrate consumption, and secreted metabolite profiles are recovered as latent variables constrained by the metabolic network stoichiometry. We resolve the core differentiability barrier of classical dFBA by reformulating the per-step linear or quadratic programme (LP/QP) as a smooth continuous ODE (the Relaxed Interior-Point ODE, R-iODE), establishing the mathematical framework for end-to-end gradient propagation through long fermentation trajectories in JAX; full gradient validation is ongoing. The result is a framework for principled inference over thousands of batch fermentations, providing a path toward model-guided medium design, cross-strain parameter transfer, and scale-up prediction from plate data.

## 1. INTRODUCTION

The design of fermentation media for probiotic production involves optimising a high-dimensional formulation space under limited experimental budget. A typical recipe contains ten or more components—glucose, citrate, amino acid pools, complex hydrolysates such as yeast extract and peptone, vitamins, and minerals—and each batch fermentation takes 32 h. At the physiological level, cells consume *substrates* from the medium, convert them through a network of metabolic reactions into new biomass, and excrete *products* such as lactate and acetate as byproducts. A predictive model must track all three: substrate depletion, biomass accumulation, and product formation.

High-throughput 96-well plate screens reduce experimental cost, but translating observed growth curves into actionable formulation decisions remains difficult (Monteiro et al., 2023). The only readily available readout in a plate screen is OD600, an optical proxy for biomass, yet medium performance depends on substrate consumption rates, secreted metabolite profiles, and intracellular fluxes that are never directly measured. Current practice either relies on empirical regression models that treat the organism as a black box, or on mechanistic models that require extensive analytical measurements to calibrate. Neither approach scales easily across strains or ingredient lots, and neither provides the uncertainty quantification needed to propagate predictions reliably to bioreactor scale.

We address this through Bayesian inference over a dynamic flux balance analysis (dFBA) model of *Lactobacillus* growth, fitting exclusively to OD600 measurements. Substrate concentrations and secreted products are *not* measured; they enter the model as latent variables whose trajectories are constrained by the metabolic network stoichiometry and inferred jointly with the kinetic parameters from the biomass signal alone. This design enables several capabilities that existing approaches do not readily provide. Once a posterior over kinetic parameters is available, medium formulations can be optimised against any user-specified objective, including ingredient cost, biomass yield, product titre, or any combination. The macronutrient weight fractions of complex hydrolysates such as yeast extract and peptone are treated as inferred parameters rather than fixed inputs, decoding opaque ingredients from growth data and making formulations more transferable across ingredient lots. Because alpha fractions are properties of the ingredient and maintenance parameters are properties of shared biochemistry, their posteriors are shared across strains and tighten from the joint dataset, reducing the calibration burden for each new organism. And because kinetic parameters describe the organism rather than the vessel, posteriors inferred from plate data carry directly to bioreactor-scale predictions without re-fitting.

Figure 1 shows the complete inference pipeline.

**Fig. 1.**
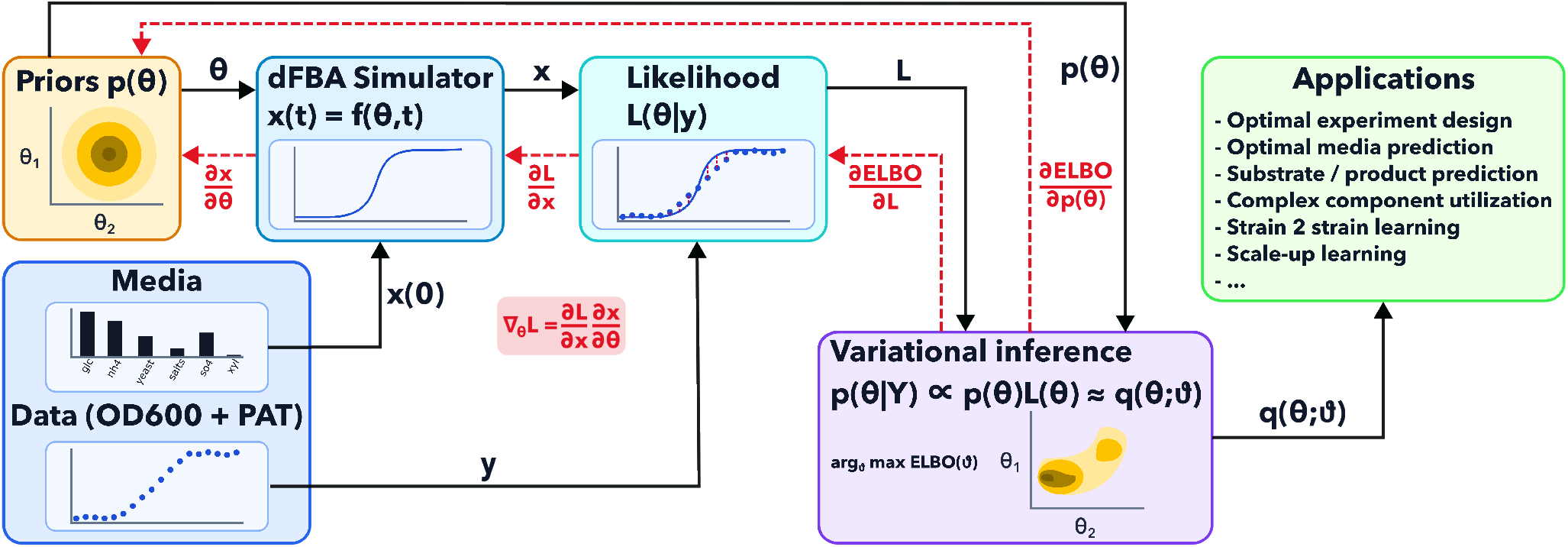
Inference pipeline. **Left:** Prior *p*(*θ*) supplies kinetic parameter samples *θ* to the dFBA simulator *x*(*t*) = *f* (*θ, t*), which is initialised from the medium composition *x*(0). **Centre:** The simulator produces substrate, biomass, and product trajectories *x*(*t*), which enter the likelihood *L*(*θ* | *y*) alongside OD600 observations *y*. The full gradient ∇_*θ*_*L* = (*∂L/∂x*)(*∂x/∂θ*) (red arrows) propagates back through the simulator via the adjoint method; *∂x/∂θ* is the key derivative enabled by the R-iODE formulation (Section 4) and is the quantity whose end-to-end validity is currently under validation (Section 8.2, Appendix G). **Right:** Variational inference fits *q*(*θ*; *ϑ*) to approximate the posterior *p*(*θ* | *Y*) by maximising the evidence lower bound (ELBO), with *∂*ELBO*/∂L* and *∂*ELBO*/∂p*(*θ*) feeding back to the likelihood and prior respectively. Inferred parameters support downstream applications including optimal media design, substrate and product prediction, complex ingredient characterisation, cross-strain transfer, and bioreactor scale-up.

**Fig. 2.**
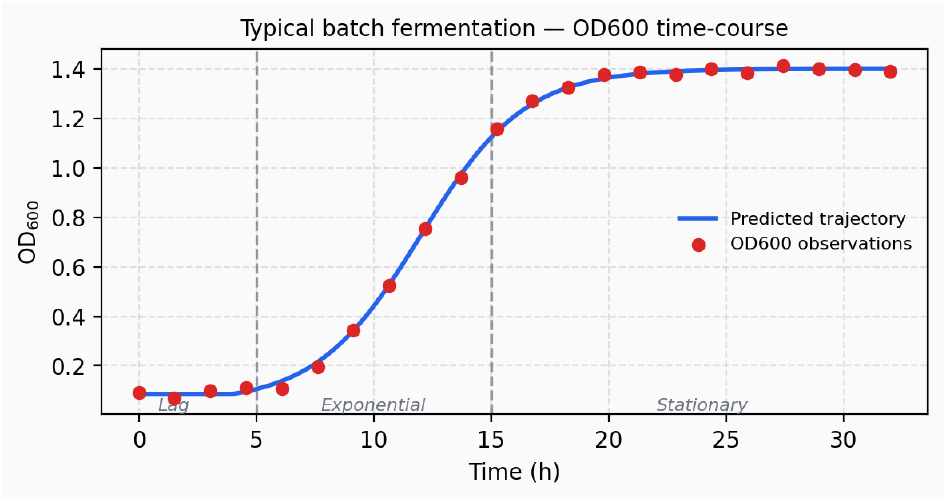
Illustrative OD600 growth curve for a *Lactobacillus* batch fermentation. The simulator must reproduce all three phases differentiably.

The central computational challenge is that evaluating the likelihood across *M* ≈ 2 100 fermentation experiments requires running *M* independent dFBA integrations, and gradient-based inference requires differentiating through all of them. Classical dFBA embeds a linear programme (LP) or quadratic programme (QP) solver inside an ODE; the solution is piecewise-linear in the constraint bounds and non-differentiable at basis changes, which occur at every ODE step as substrate concentrations evolve. This paper describes how we resolve the differentiability problem and build a simulator suitable for variational inference at this scale.

### Relation to existing work

Dynamic flux balance analysis was introduced by Mahadevan et al. (2002) and has since become widely adopted by experimental practitioners for modelling batch fermentations (Lewis et al., 2012). The need for differentiability in metabolic simulators has been recognised in several lines of work. Amos and Kolter (2017) showed that the KKT conditions of a QP can be differentiated implicitly (OptNet), yielding valid gradients at a fixed optimal basis. However, in dFBA the active set changes at every ODE step as substrate concentrations evolve; OptNet’s implicit derivatives are discontinuous across these basis changes and therefore unsuitable as a drop-in replacement. Entropy-regularised LP relaxations produce a globally smooth approximate solution, but the approximation error accumulates over long trajectories and distorts the inferred kinetics—a particularly serious problem when the goal is parameter inference rather than control. Scott et al. (2018) proposed a qualitatively different approach: rather than differentiating through a solver, one can embed the KKT stationarity conditions of a fixed-barrier QP directly as an ODE for the flux coordinates, yielding a smooth dynamical system whose right-hand side requires only a linear solve per step. Scott demonstrates this Relaxed Interior-Point ODE (R-iODE) formulation for simulation, and suggests that its differentiability makes it suitable for parameter estimation—but stops short of actually doing so. To our knowledge, the present work is the first to carry out gradient-based Bayesian inference through an R-iODE, integrating it into a full variational inference pipeline over thousands of experiments.

Our implementation departs from Scott et al. (2018) in several respects that prove essential for making the approach computationally tractable at this scale. Scott works in the full reaction flux space *v* ∈ ℝ^|ℛ|^ and must track the Lagrange multiplier for the stoichiometric equality *Sv* = 0 as a differential state alongside *v*. We instead project onto the null space of *S*, replacing *v* with coordinates *u* ∈ ℝ^*r*^, *r* = −rank(*S*), so that *Sv* = 0 is satisfied automatically for any *u*. This removes the equality-constraint multiplier from the ODE state entirely—it is uniquely determined algebraically by the complementarity condition—and reduces the ODE dimension from ~50 to ~15 for our *L. rhamnosus* model. Smaller state means lower stiffness, faster integration, and a smaller Jacobian for the adjoint method. We further introduce medium-specific trimming to maintain strict interior feasibility, and three smooth gating functions—near-zero Monod tapering, an ATP maintenance softmin, and a biomass objective sigmoid—to handle the biological boundaries that arise in batch fermentation and that are absent from the chemical engineering setting of Scott et al. (2018).

A related and concurrent line of work applies KKT-based reformulation of the dFBA LP to model predictive control of bioprocesses (Nakama and Jäschke, 2022b,a; de Oliveira and Jäschke, 2025; Gotsmy et al., 2024). These approaches replace the embedded LP with its KKT conditions and solve the resulting nonlinear programme using a general-purpose interior-point solver (IPOPT) that drives *µ* → 0, targeting the exact LP optimum for offline trajectory optimisation. The R-iODE differs in two respects: *µ* is fixed permanently at a positive value, keeping the solution on the central path and ensuring global smoothness; and the optimality conditions are embedded as a continuous ODE rather than an NLP constraint, which is the property that enables adjoint-method differentiation through the full integration trajectory. The two approaches are thus complementary—KKT-based NLP for control problems where exactness matters, R-iODE for inference problems where differentiability is the primary requirement.

Beyond the simulator itself, the probabilistic model introduced here addresses a gap in the existing dFBA inference literature. Most dFBA inference studies assume fully characterised, pure-component media. Industrial fermentation media—yeast extract, peptones, meat extract—contain macronutrients in unknown proportions that vary by supplier and batch. We introduce *alpha fractions* as learnable parameters encoding the weight fractions of macronutrient classes (carbohydrates, protein) in each complex ingredient, inferred jointly with the kinetics. To our knowledge this has not previously been done in a probabilistic dFBA framework.

A complementary consideration motivates the Bayesian treatment over a point estimate: inference should distinguish parameters that are shared across strains from those that are strain-specific. Alpha fractions and maintenance requirements are properties of the ingredient or of conserved biochemistry; their posteriors tighten rapidly from the joint dataset and transfer directly to a new strain without re-estimation. Strain-specific kinetic parameters (Monod constants, cybernetic affinities) require fresh experiments, but the shared parameters constrain the feasible region and reduce the experimental burden considerably.

Finally, we note that a point estimate 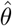 collapses all uncertainty to a single value and cannot propagate that uncertainty to downstream decisions—which medium to test next, which experiment would be most informative, whether a scale-up prediction is reliable enough to act on. The full posterior *p*(*θ*| 𝒟) supports all of these through predictive intervals and information-theoretic experimental design criteria.

#### 1.1 Comparison with regression and neural ODE baselines

Three broad modelling strategies are applicable to fermentation growth curve data, and it is instructive to contrast their representational scope.

##### Regression on summary statistics

The simplest approach fits a parametric growth model—typically a Gompertz curve—to each trajectory and regresses the resulting summary statistics (maximum OD, growth rate, lag time) against the input medium composition. This requires assuming a fixed functional form for the growth trajectory, which fails for biphasic or asymmetric profiles. More fundamentally, the approach operates entirely in the space of the observable (biomass); substrates and products are not represented and cannot be predicted.

##### Neural ODEs

A neural ODE (Chen et al., 2018) learns the differential equation governing biomass dynamics directly from data, removing the Gompertz assumption and recovering the flexibility to fit complex trajectories. However, if only biomass is observed, the neural ODE state is biomass; substrates and products remain absent because there is no structure in the model that would couple them to the observable signal.

##### Grey-box dFBA simulator (this work)

Our simulator models the same complex growth dynamics but maintains the full extracellular state *c*^*e*^ = (*c*^subs^, *c*^prod^, *c*^*bm*^)^*T*^. The metabolic network stoichiometry constrains how substrates, products, and biomass co-evolve: substrate depletion must be consistent with the observed biomass gain, and product excretion must close the carbon balance. Substrate and product trajectories are therefore identifiable as latent variables from OD600 alone, without requiring direct measurement. This is the representational property that makes the approach relevant to the four objectives above.

## 2. DYNAMIC FLUX BALANCE ANALYSIS

A stoichiometric model represents metabolism as *S* ∈ ℝ^| ℳ|×|ℛ|^, where *S*_*ij*_ is the stoichiometric coefficient of metabolite *i* in reaction *j*. At quasi-steady state, *Sv* = 0. Flux balance analysis (FBA) (Orth et al., 2010) predicts growth by maximising biomass flux subject to this equality and bound constraints *A*^*v*^*v* ≤ *b*. Parsimonious FBA merges with a quadratic regulariser to form a QP:

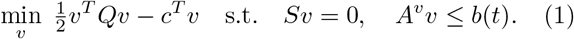

### 2.1 The extracellular ODE

dFBA (Mahadevan et al., 2002) tracks extracellular concentrations *c*^*e*^ = (*c*^subs^, *c*^prod^, *c*^*bm*^)^*T*^ :

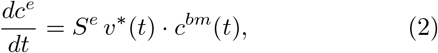

where *S*^*e*^ is the exchange stoichiometry submatrix and *v*^*^(*t*) solves (1). The time-varying import bounds *b*^subs^(*t*) are given by Monod kinetics,

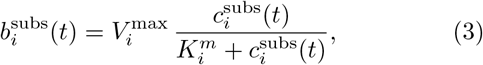

which couple the extracellular state back to the QP through the bound vector *b*(*t*).

The solution *v*^*^(*t*) is piecewise-linear in *b*(*t*) and therefore non-differentiable at basis changes of the LP/QP. Because *c*^*e*^(*t*) evolves continuously, multiple such activeset transitions occur along a typical trajectory, and the gradient *∂v*^*^*/∂b* is undefined at each one. By the chain rule, *∂v*^*^*/∂θ* inherits this non-differentiability, blocking gradient computation for any downstream loss.

## 3. NULL-SPACE REDUCTION AND MEDIUM-SPECIFIC TRIMMING

### 3.1 Eliminating equality constraints

Let *N* ∈ ℝ^|ℛ|×*r*^ span ker(*S*), *r* = |ℛ| − rank(*S*). Every feasible *v* = *Nu* satisfies *Sv* = 0 automatically. The QP (1) becomes:

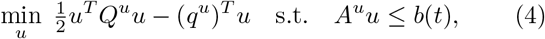

with *Q*^*u*^ = *N*^*T*^ *QN*, *q*^*u*^ = *N*^*T*^ *c, A*^*u*^ = *A*^*v*^*N*. For our curated *L. rhamnosus* model this reduces from |ℛ| ≈ 50 to *r* ≈ 15 null-space coordinates. Flux variability analysis (FVA) pre-removes constraints in *A*^*v*^ that are never active at any feasible operating point.

### 3.2 Medium-specific polytope trimming

The ~ 2 100 fermentation experiments decompose into 54 unique medium compositions when classified by their pattern of non-zero components (see Appendix A for a full description of the dataset and how this count is derived). This reduction is exploited to precompute a separate trimmed polytope for each unique medium composition, amortising the cost across the many experiments that share the same pattern.

The log-barrier formulation requires the slack variables *s* = *b*(*t*) − *A*^*u*^*u* to be strictly positive throughout the integration. A substrate with zero initial concentration has a Monod upper bound of zero at *t* = 0 and remains at zero for all *t*: its import reaction is permanently inactive. If such a reaction is retained in the null-space coordinate system, the corresponding import constraint takes the form 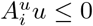 with an initial slack of zero. The polytope then has no interior point satisfying all constraints with *s >* 0, and the log-barrier diverges immediately.

We therefore trim each medium-specific polytope before integration by removing all reactions whose maximum feasible flux under the prior is zero, computed via a per-reaction linear programme over the base polytope with medium-specific upper bounds. Formally, reaction *j* is removed if

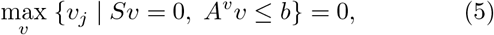

where the import upper bounds in *b* are set to zero for all absent substrates. Removing zero-flux columns from *S* may leave metabolite rows with no remaining non-zero entry; these rows are eliminated iteratively until *S* has no empty rows. Redundant inequality constraints are subsequently removed, and priority constraints (Monod import bounds and the NGAM lower bound) are preserved unconditionally.

The resulting trimmed system has a medium-specific stoichiometric matrix *S*^(*k*)^ ∈ ℝ^|M(*k*) |×|ℛ(*k*) |^ and null-space basis *N* ^(*k*)^, with |ℛ| ^(*k*)^ ≪ |ℛ| for media containing only a subset of substrates. The polytope of the trimmed system has a well-defined interior, and the log-barrier is well-conditioned from *t* = 0.

A consequence of this trimming is that the Monod parameters 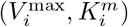 for substrate *i* are identifiable only from experiments in which substrate *i* is present. The ~85 unique medium patterns span different subsets of substrates, so different parameter subsets are constrained by different experiment subsets. This structure is captured naturally by the Bayesian framework: parameters for substrates that appear in few media have wider posteriors, and the shared prior regularises them towards biologically plausible values.

## 4. THE R-IODE FORMULATION

### 4.1 Log-barrier QP at fixed µ

We replace inequality constraints with a log-barrier:

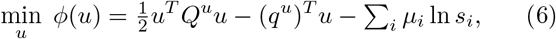

with *s* = *b*(*t*) − *A*^*u*^*u >* 0. In a standard interior-point solver *µ* → 0; the key insight of Scott et al. (2018) is to *fix µ >* 0 *permanently*. The barrier-optimal solution *u*^*^(*t*) then tracks the central path of the QP as *b*(*t*) evolves—never reaching the constraint boundary, always smooth and differentiable.

### 4.2 KKT conditions as an ODE

First-order optimality of (6) gives stationarity *Q*^*u*^*u* − *q*^*u*^ + (*A*^*u*^)^*T*^ *λ* = 0 and complementarity *s*_*i*_*λ*_*i*_ = *µ*_*i*_. The multipliers are recovered algebraically: *λ*_*i*_ = *µ*_*i*_*/s*_*i*_.

Differentiating both conditions with respect to time and collecting terms yields a block linear system solved at every ODE step:

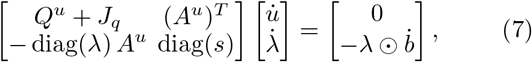

where 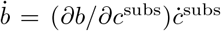 from Monod kinetics and *J*_*q*_ is the Jacobian of any state-dependent modification to *q*^*u*^ (see Section 5). We retain only the 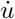 block; *λ* is recomputed algebraically at each step.

### 4.3 Full ODE

The complete state is *x* = [*c*^*e*^, *u*]^*T*^ :

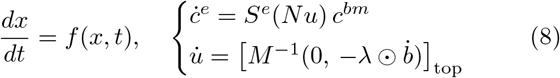

One linear solve per ODE step; no LP or QP at runtime; no active-set switches. The initial condition *u*^0^ is found by solving (4) once at *t* = 0. Gradients flow through the full trajectory via the adjoint method (Chen et al., 2018; Kidger, 2022). Figure 4 illustrates the geometry of a single ODE step: the left panel shows the feasible polytope at *t* = 0 with abundant substrates, where the log-barrier shading keeps the solution (green dot) well away from every constraint face; the right panel shows the same polytope late in the batch, after Monod upper bounds have contracted inward— the R-iODE has followed the central path continuously without any active-set event.

**Fig. 3.**
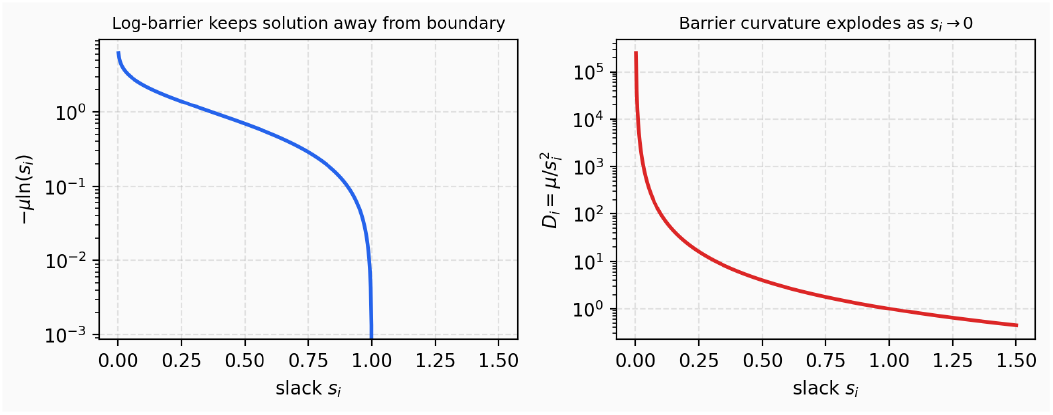
Log-barrier −*µ* ln *s*_*i*_ (left) and barrier curvature 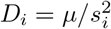 (right). The curvature diverges as *s*_*i*_ → 0, motivating the gating functions in Section 5.

**Fig. 4.**
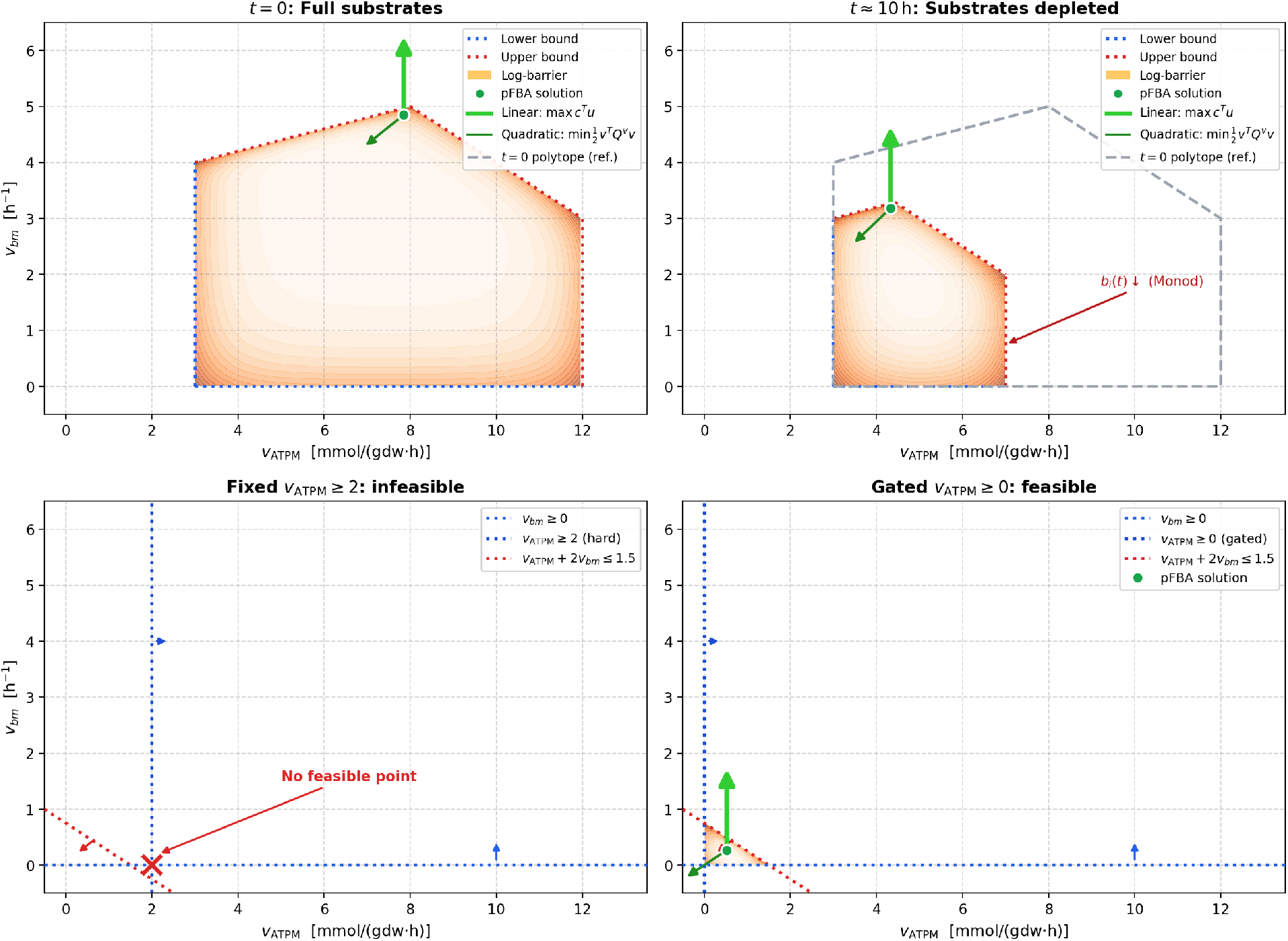
Flux polytope projected onto the *v*_ATPM_–*v*_*bm*_ plane. *Top row* : polytope evolution along a batch—full substrates at *t* = 0 (top-left) and after depletion at *t* ≈ 10 h (top-right); the R-iODE tracks the central path continuously as Monod bounds contract. *Bottom row* : ATPM feasibility—the hard NGAM lower bound leaves no feasible point late in the batch (bottom-left); the softmin gate lowers the effective bound to zero and restores a small feasible region (bottom-right).

#### A note on the fixed-barrier optimum

By fixing *µ >* 0 permanently, the R-iODE tracks the *central path* of the QP rather than its exact LP optimum. Kinetic parameters inferred under this formulation are therefore slightly biased relative to parameters that would be identified under classical dFBA with an exact solver. We argue this bias is inconsequential for two reasons. First, *µ* is fixed at a small value (*µ* ≲ 10^−5^), so the central-path solution is very close to the constraint boundary. Second, and more fundamentally, dFBA is itself an imperfect model of cellular metabolism: the maximum-biomass objective, fixed enzyme capacity bounds, and quasi-steady-state flux assumption each introduce structural modelling errors. These structural errors exceed the barrier bias by orders of magnitude. The inferred posterior is therefore still approximately correct and useful for the intended downstream applications—medium design, scale-up prediction, and strain transfer—even though it is not identical to the posterior one would obtain under an exact LP solver.

## 5. GATING FUNCTIONS

The R-iODE as derived in Section 5 assumes that all slacks *s*_*i*_ *>* 0 are bounded away from zero throughout the integration. Three smooth modifications are required to maintain this condition under the biological dynamics of batch fermentation; Figure 5 shows each one.

**Fig. 5.**
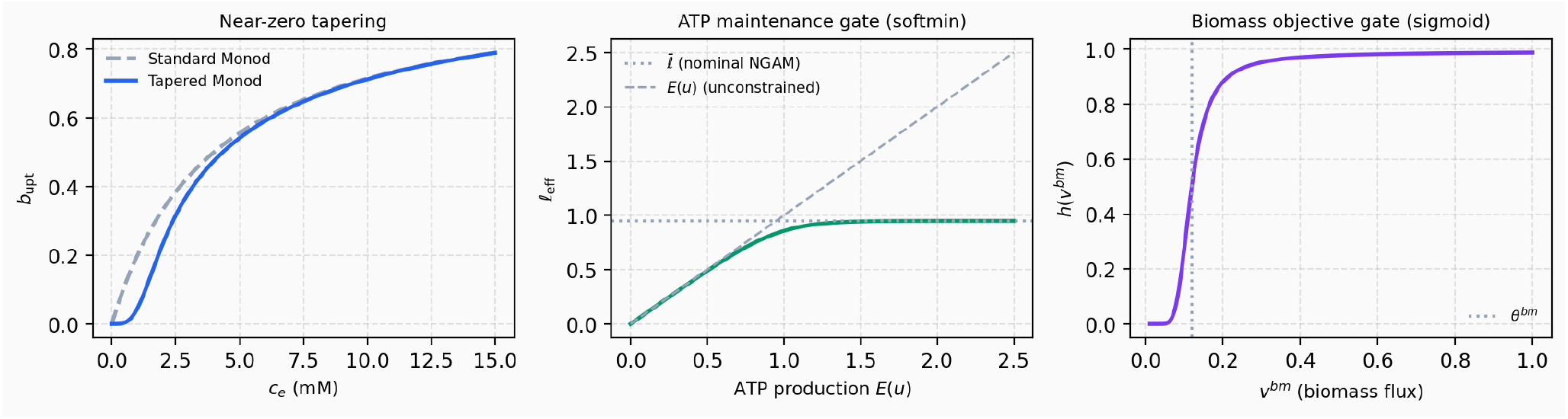
The three smooth gating functions: near-zero Monod tapering (*left*), ATP maintenance softmin (*centre*), and biomass objective sigmoid (*right*).

### 5.1 Monod near-zero tapering

Standard Monod kinetics have a non-vanishing derivative at *c*_*i*_ = 0. We multiply the Monod bound by a smooth tapering factor:

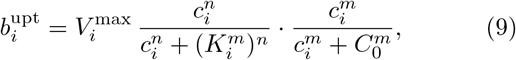

with *m* = 3, *C*_0_ = 1.0 mM. As shown in the left panel of Figure 5, the taper factor approaches zero as *c*_*i*_ → 0, ensuring that the import bound and the corresponding slack collapse continuously rather than abruptly.

### 5.2 ATP maintenance gate (softmin)

Non-growth-associated maintenance (NGAM) imposes a strictly positive lower bound 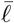 on the ATP production flux. _*i*_

As substrates are consumed over the course of a batch, the feasible polytope can shrink until no flux vector satisfies this bound simultaneously with the import constraints, rendering the QP infeasible. To maintain a non-empty feasible region throughout the integration, we replace the hard NGAM bound with a smooth lower bound:

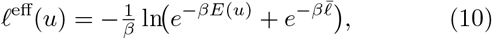

Where 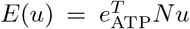. When 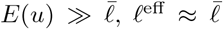. As *E*(*u*) → 0, the bound relaxes to zero, keeping the polytope non-empty throughout the batch (Figure 5, centre panel). The bottom row of Figure 4 shows the polytope geometry: the bottom-left panel illustrates the late-batch scenario where Monod upper bounds shrink the feasible region below the NGAM corner so no feasible point exists under the hard bound; the bottom-right panel shows how the softmin gate lowers the effective lower bound and restores a small feasible region near the origin.

### 5.3 Biomass objective gate (sigmoid)

Near batch end, the linear objective in (6) produces large 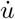 for small changes in *b*(*t*), causing numerical blow-up. We gate the linear cost vector down as achievable biomass flux declines:

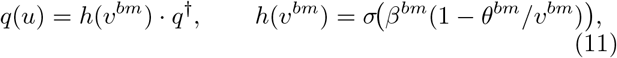

Where 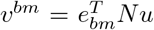. As *v*^*bm*^→ 0, *h* → 0, the quadratic cost takes over, and *u* → 0 smoothly (Figure 5, right panel). The Jacobian *J*_*q*_ = *∂q/∂u* feeds into (7).

The scalar *q*^†^ is set at initialisation to the *minimum* value for which the *µ*-relaxed QP achieves at least 99.5% of its plateau biomass flux—guaranteeing that the quadratic penalty dominates the linear driving force while leaving maximum dynamic range for the gate. Specifically, *q*^†^ is found via a geometric search to locate the plateau followed by bisection to target the 99.5% threshold (at fixed *b* and *µ*). When *h*(*t*) *<* 1, the effective objective *h*(*t*) · *q*^†^ drops proportionally, so the gate only activates when the achievable biomass flux itself has declined, rather than suppressing growth prematurely.

## 6. CYBERNETIC CONTROL

*Lactobacillus* preferentially consumes the most energetically favourable carbon source first—the classic diauxic growth pattern (Monod, 1942). We model resource allocation with a softmax weighting:

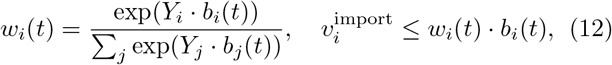

where *Y*_*i*_ is the ATP yield of substrate *i*.

A numerical difficulty arises from the exponential sensitivity of the softmax: when a high-yield substrate such as glucose is abundant, its logit can exceed those of secondary substrates by tens of nats, suppressing their allocations to values near machine zero. The effective import bounds for those substrates collapse accordingly, the corresponding slacks approach zero, and the barrier Hessian 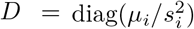 diverges. To prevent this, we apply a uniform mixing floor:

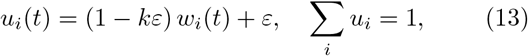

with *k* substrates and *ε* = 0.01, guaranteeing a minimum allocation *ε* regardless of logit magnitude. Figure 6 demonstrates the effect on a two-substrate batch: the left panel traces substrate concentrations, with glucose consumed first in accordance with the diauxic preference; the centre panel shows that under pure softmax the effective import bound *u*_*i*_*b*_*i*_ for the secondary substrate collapses to near machine zero while glucose is present, causing the barrier Hessian 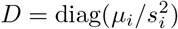 to diverge; the right panel confirms that the mixing floor with *ε* = 0.01 maintains a small but non-zero effective bound throughout the batch, keeping the Hessian well-conditioned.

**Fig. 6.**
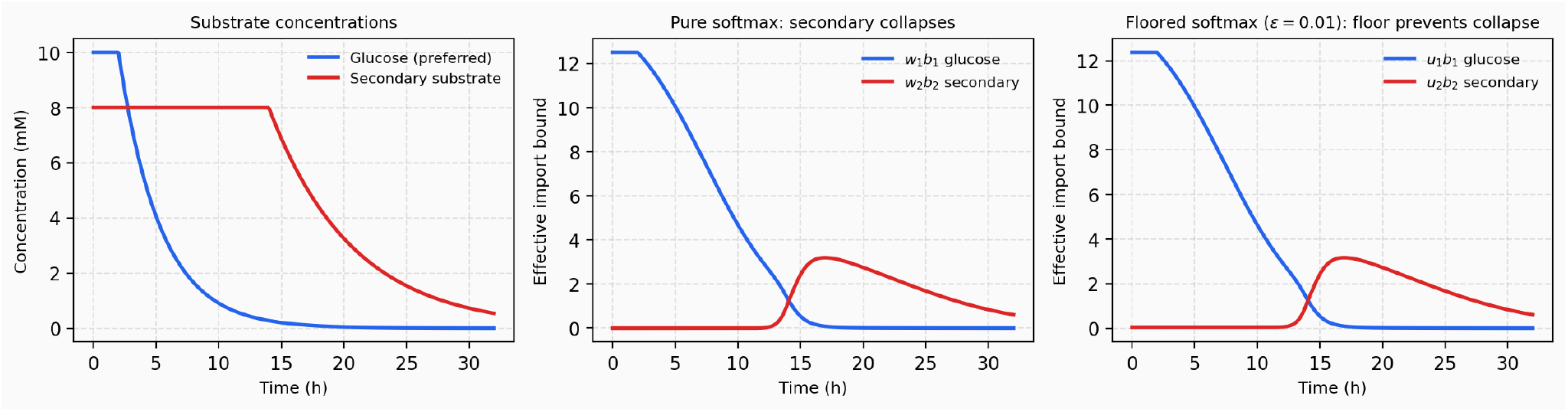
Cybernetic resource allocation: substrate concentrations (*left*), effective import bounds *u*_*i*_*b*_*i*_ under pure softmax (*centre*), and with uniform mixing floor *ε* = 0.01 (*right*).

## 7. PROBABILISTIC MODEL

### 7.1 Parameters and prior structure

The parameter vector *θ* contains approximately 60 entries per strain, organised into five groups: Monod kinetics 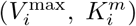, stoichiometric coefficients, alpha fractions encoding the macronutrient composition of complex ingredients, and an OD600 observation model. Full prior specifications are given in Appendix B.

### 7.2 Likelihood

The likelihood factorises over experiments and time points:

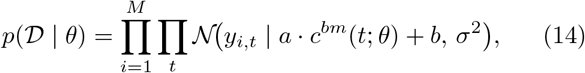

where *c*^*bm*^(*t*; *θ*) is the biomass trajectory from integrating (8). Because the simulator produces full substrate and product trajectories alongside biomass, the likelihood is easily extended to any measurement that observes the simulated state: end-point substrate or metabolite assays (e.g. HPLC), or dense time-course signals from inline process analytical technology (PAT) such as Raman spectroscopy or mass spectrometry. Each additional signal type enters as an independent factor in the likelihood with no change to the simulator or inference architecture. The posterior is approximated via a neural spline flow trained by maximising the ELBO; details are given in Appendix C.

## 8. IMPLEMENTATION

The simulator is implemented in JAX (Bradbury, 2018) with 64-bit floating point and integrated with `diffrax` (Kidger, 2022) using a stiff implicit solver (Kvaerno-5 SDIRK) with adaptive step-size control. The metabolic model is loaded and manipulated with COBRApy.

Static quantities (null-space basis *N, A*^*u*^, *Q*^*u*^, FVA-trimmed constraints) are computed once at initialisation per unique experimental condition. Dynamic quantities (Monod bounds, slacks, multipliers, gating values) are recomputed at each ODE step as pure functions of the current state, which is required for JIT compatibility.

### 8.1 JIT compilation

Because all dynamic quantities are expressed as pure JAX functions with no Python side-effects, the full ODE right-hand side—including the block linear solve (7)— can be traced and compiled by XLA. The first call to `simulate_ode` incurs a one-time tracing and compilation cost of approximately 2 s; all subsequent calls with the same experiment structure run the compiled XLA kernel in approximately 0.03 s, a ~60× speedup. This makes mini-batch gradient steps feasible: a batch of 16 experiments can be simulated in under 1 s on a single GPU.

### 8.2 Gradients and the adjoint method

The adjoint method (Chen et al., 2018) differentiates through the ODE trajectory without storing intermediate activations, making gradient computation memory-efficient even over long integration windows. `diffrax` implements a recursive-checkpoint variant that balances memory and recomputation cost (Kidger, 2022). The block linear solve (7), all three gating functions, and the Monod kinetics are JAX-autodiff-compatible, so gradients flow end-to-end from the biomass trajectory to the kinetic parameters— specifically, *∂x/∂θ* in Figure 1 is computed via the adjoint pass through the ODE. In the current implementation, the forward ODE integration is stable for the majority of prior draws (Appendix F), but the backward adjoint pass through the KKT linear system (7) produces non-finite values for all tested (experiment, seed) pairs. This is a numerical issue in the current implementation, not a fundamental limitation of the R-iODE formulation: the forward pass is fully differentiable by construction and the KKT block solve (7) has well-defined analytical gradients. The root cause is that the KKT matrix *M* in (7) can become ill-conditioned during backward integration, causing the linear solver to return NaN or Inf. Resolving this—for instance by regularising *M* during the adjoint pass, or by applying the `stop_gradient` operator to the dual variables *λ* in the backward pass—is the active engineering task that gates variational inference (see Appendix G).

## 9. SIMULATION RESULTS AND COMPLETION RATE

Figure 7 shows a representative successful integration: substrate concentrations decline monotonically, biomass accumulates, and the import flux trajectories respect both the Monod kinetic bounds and the cybernetic allocation. The KKT condition number (Appendix D) remains below 10^12^ throughout, and all slack variables stay bounded away from zero.

**Fig. 7.**
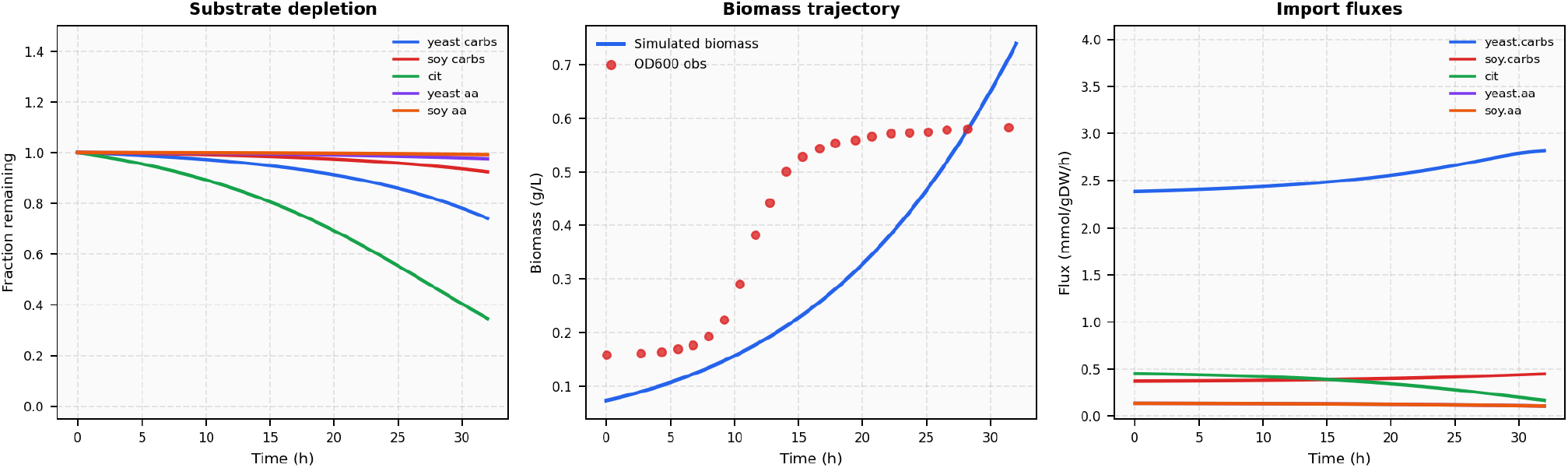
Representative successful simulation for a *L. rhamnosus* batch (exp. 1974, prior draw seed 0). Substrate concentrations decline as biomass accumulates; import fluxes track Monod kinetics and cybernetic allocation. Diagnostic plots (slack variables and KKT condition number) for this run are given in Appendix D.

### 9.1 Completion rate

Not every prior draw integrates successfully to the batch end-time *t*_1_ = 32 h. We quantified this with two sweeps:

**Sweep A** held experiment 1974 fixed and varied the parameter seed (30 draws); **Sweep B** held the seed fixed and varied the experiment index across the full dataset (30 experiments, evenly spaced). Full results are shown in Appendix F. Completion is defined as reaching *t* ≥ 30 h. Sweep A reached 25/30 (83%); Sweep B reached 29/30 (97%). The dominant failure mode is ODE step-size collapse driven by a combination of substrate depletion, ATP maintenance infeasibility, and cybernetic softmax overflow, which cause the KKT system (7) to become ill-conditioned. The gating functions in Section 5 address all three mechanisms; improving completion rate over the full prior support is an active objective.

Completing the integration to *t*_1_ is a prerequisite for likelihood evaluation: a truncated trajectory has no final biomass value and therefore contributes no gradient to the ELBO (C.1). Experiments whose simulations fail are dropped from the current training batch, which reduces effective batch size and slows convergence. Raising the completion rate is therefore the primary remaining engineering challenge before full variational inference can be run.

## 10. DISCUSSION AND CONCLUSION

We have described a differentiable dFBA simulator that replaces per-step LP/QP solves with a smooth ODE derived from the KKT conditions of a fixed-barrier interior-point programme. Null-space reduction compresses the ODE dimension by 3–4 ×; medium-specific trimming maintains strict interior feasibility; three gating functions extend the formulation to the full range of batch dynamics. The result is a JAX-differentiable system compatible with adjoint-method differentiation, making variational inference over ~ 2 100 fermentation experiments tractable.

The principal methodological contributions are: (i) the first end-to-end gradient-based Bayesian inference pipeline through an R-iODE, extending Scott et al. (2018) from simulation to parameter estimation; (ii) a null-space reduction that reduces the ODE state dimension by 3–4× relative to full reaction-space formulations; (iii) medium-specific polytope trimming that maintains strict interior feasibility across diverse media compositions; and (iv) smooth gating functions that handle the full range of batch dynamics, including near-zero substrate depletion, ATP maintenance switching, and biomass collapse.

The kinetic parameters *θ* are properties of the organism and its metabolic network, not of the fermentation vessel. A posterior distribution obtained from 96-well plate data is therefore directly applicable to bioreactor-scale predictions: the hydrodynamic and mass transfer environment changes between scales, but the enzymatic kinetics do not. This is the principal practical motivation for the Bayesian treatment over a point estimate.

The alpha-fraction posteriors carry additional scientific value independent of scale-up. They quantify the macronutrient weight fractions—carbohydrates, protein—of each complex hydrolysate as experienced by the organism: a quantity that cannot be measured directly and is not reported in supplier specifications. Because alpha fractions are properties of the ingredient rather than the organism, their posteriors are shared across strains and tighten rapidly from the joint ×2 100-experiment dataset, reducing the experimental burden for each new strain characterisation.

The present model does not account for pH dynamics and product inhibition, which become relevant in prolonged fermentations, nor for the lag phase observed when inoculum quality varies. These are identified as natural extensions of the current framework.

The design choices made here are not inherently specific to *Lactobacillus* or probiotic production. Any organism with a sufficiently curated genome-scale metabolic model and accessible batch OD600 data is a potential candidate for this inference pipeline. Extensions to fed-batch or continuous culture would require modifications to the extracellular ODE but not to the core R-iODE or inference machinery. The framework is also complementary to targeted metabolomics: posterior distributions over latent substrate and product trajectories identify which additional measurements would most reduce parameter uncertainty, supporting more efficient experimental design campaigns.

The differentiable simulator described here is complementary to KKT-based dFBA formulations used in model predictive control (Nakama and Jäschke, 2022b; de Oliveira and Jäschke, 2025; Gotsmy et al., 2024): once posterior distributions over kinetic parameters are available from the inference pipeline, they can be propagated forward as uncertainty scenarios in an economic MPC framework, connecting the two lines of work into an end-to-end decision pipeline from plate screening to process control.

## Appendix A. EXPERIMENTAL DATASET

### Medium components

Each experiment is initialised from a medium specified as concentrations (g L^−1^) of up to 18 ingredients, grouped by function:

- **Complex hydrolysates**: yeast extract, meat extract, soy peptone, pea peptone.
- **Carbon sources**: glucose, sucrose. **Nitrogen/carbon salts**: ammonium hydrogen citrate, sodium acetate, ammonium acetate.
- **Mineral salts**: magnesium sulfate, potassium phosphate dibasic, manganese sulfate monohydrate, sodium chloride.
- **Cysteine sources**: cysteine, cysteine HCl monohydrate.
- **Modifiers**: Tween 80, ascorbic acid, glutathione.

### From ingredients to model inputs

Ionic salts dissociate in solution, so each salt is mapped to its constituent ions: magnesium sulfate contributes Mg^2+^ and 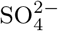; potassium phosphate dibasic contributes K^+^ and inorganic phosphate; and so on for the remaining salts. We do not model the transport dynamics of the counterions (K^+^, Mg^2+^, Mn^2+^, Na^+^, Cl^−^) because they are not rate-limiting under the conditions studied. As a result, two media that differ only in which mineral salts are present but supply the same carbon and nitrogen sources are treated as the same composition by the model.

Complex hydrolysates (yeast extract, peptones) are parameterised by alpha fractions that encode the macronutrient weight fractions of carbohydrates, protein, and nucleotides; these are inferred jointly with the kinetic parameters (Section 7).

### Dataset size

We use two terms throughout. A *composition* is the set of ingredients that are present (non-zero) in a medium, regardless of their amounts—for example, a medium

containing glucose and yeast extract at any concentration is one composition. A *concentration profile* is a specific set of amounts for a given composition; multiple experiments can share the same composition but differ in the concentration levels of their ingredients, each contributing an independent OD600 trajectory.

The *L. rhamnosus* dataset contains 2 121 experiments after duplicate removal. At the metabolic level these span 140 distinct compositions. Excluding ion transport reactions (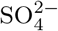, inorganic phosphate) causes 84 of these 140 compositions to become duplicates of others and be merged, reducing the count from 140 to 56: media that differed only in mineral salt amounts but shared the same active carbon and nitrogen sources are now treated identically. A further feasibility screen using flux balance analysis removes 2 additional compositions for which biomass production is not achievable under any flux allocation, leaving **54 feasible compositions** in total. Each composition requires a distinct ODE system; the kinetic parameters *θ* are shared across all of them. Figure A.1 shows how the 2 121 experiments are distributed across these 54 compositions.

**Fig. A.1.**
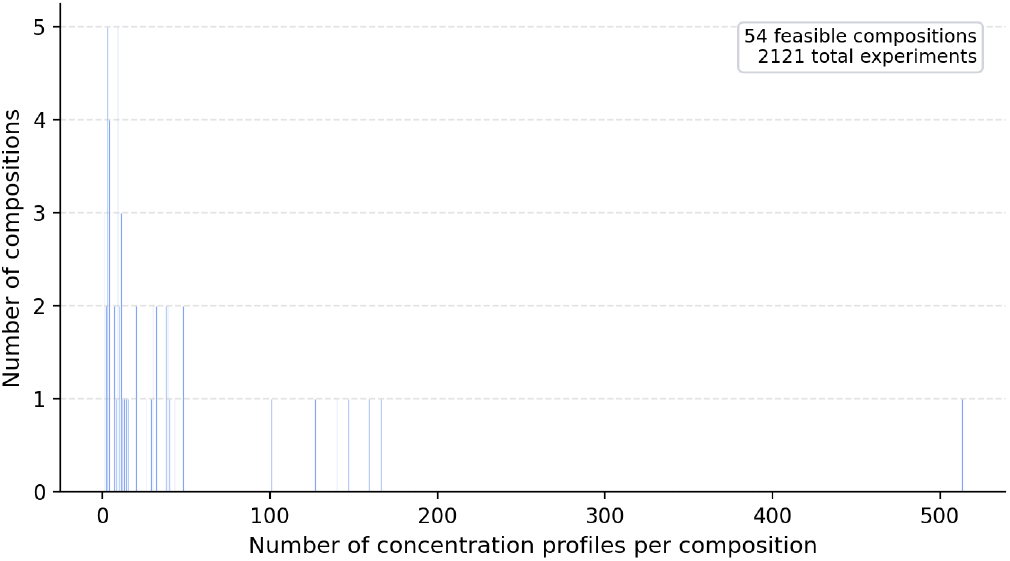
Distribution of concentration profile counts across the 54 feasible *L. rhamnosus* medium compositions. Each bar represents the number of compositions with a given number of replicate concentration profiles. The majority of compositions are covered by fewer than 50 profiles; a small number of high-frequency compositions account for a disproportionate share of the 2 121 total experiments.

## Appendix B PRIOR SPECIFICATION

The parameter vector *θ* contains approximately 60 entries per strain, organised into five groups with distinct prior structure.

### Monod kinetics

Each import reaction carries 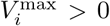 and 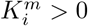. We use OffsetLogNormal priors with a hard lower bound *δ* to keep parameters away from zero. Mean values are informed by measured *L. rhamnosus* saturation constants (Polak-Berecka, 2010) and amino-acid symporter costs (Poolman, 1993; Strobel, 1989).

**Fig. B.1.**
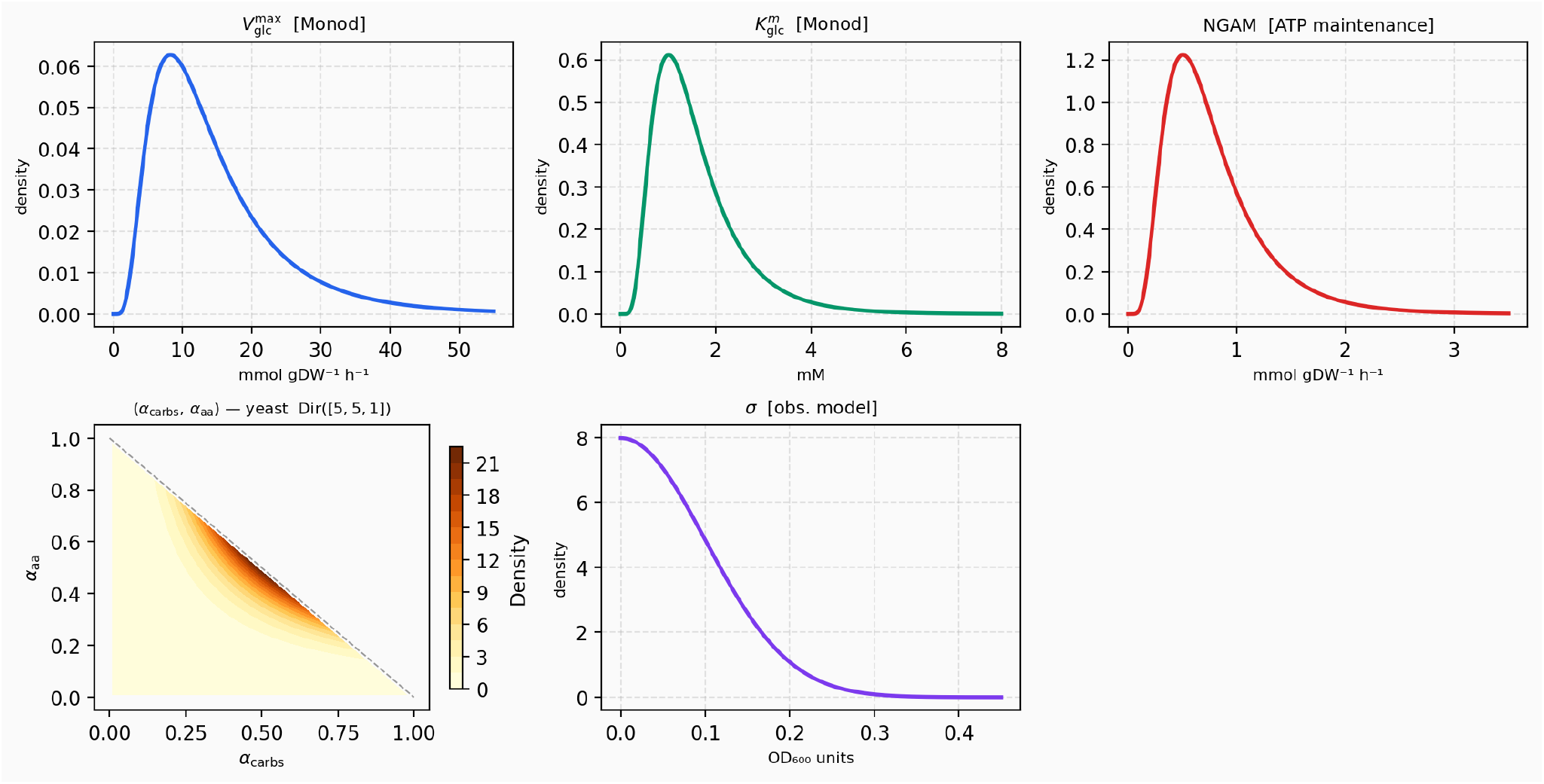
Prior distributions over the five parameter groups. OffsetLogNormal priors bound positive quantities away from zero. Dirichlet priors on alpha fractions encode the simplex constraint.

### Stoichiometric coefficients

Unknown entries in *S*—ATP cost of amino acid import, growth-associated maintenance (GAM)—follow OffsetLogNormal priors.

### Alpha fraction

Fractions 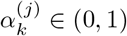 split each complex ingredient *k* into metabolite categories *j*:

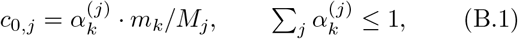

where *m*_*k*_ is the component concentration and *M*_*j*_ a representative molar mass. These are shared across strains (yeast-extract composition does not depend on which organism consumes it) and are inferred jointly from all three *Lactobacillus* strains. Dirichlet priors encode the simplex constraint.

### Observation model

OD_600_ = *a* · *c*^*bm*^ + *b* + *ε, ε* ~ 𝒩 (0, *σ*^2^), with *a, b* following tight OffsetLogNormal priors.

## APPENDIX C VARIATIONAL INFERENCE

The posterior *p*(*θ* | 𝒟) ∝ *p*(*θ*| 𝒟) *p*(*θ*) is intractable in 60 dimensions. We fit an approximate distribution *q*_*ϕ*_(*θ*) by maximising the ELBO:

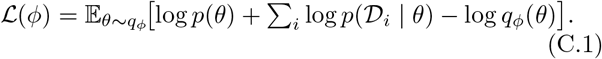

We use a **neural spline flow** (Durkan et al., 2019) as *q*_*ϕ*_: a composition of invertible transformations mapping *z*_0_ ~ 𝒩 (0, *I*) to *θ*, where each univariate bijection is a monotone rational-quadratic spline. Splines represent the skewed, bounded posteriors over kinetic parameters more faithfully per layer than affine transforms (Papamakarios et al., 2021). The flow is composed with a fixed bijection *g* (softplus for positive parameters, isometric log-ratio for Dirichlet variables) so that samples automatically lie in the constrained support.

Gradients are estimated via the reparameterisation trick (Rezende and Mohamed, 2015) on mini-batches of *B* = 4–16 experiments. The gradient path ∇_*ϕ*_ ℒ requires differentiating through the ODE simulation—made possible by the adjoint method (diffrax) and the differentiable block solve (7).

## Appendix D SIMULATION DIAGNOSTICS — SUCCESSFUL RUN

Figure D.1 shows nine diagnostic time series for the successful simulation in Section 9 (exp. 1974, seed 0).

**Fig. D.1.**
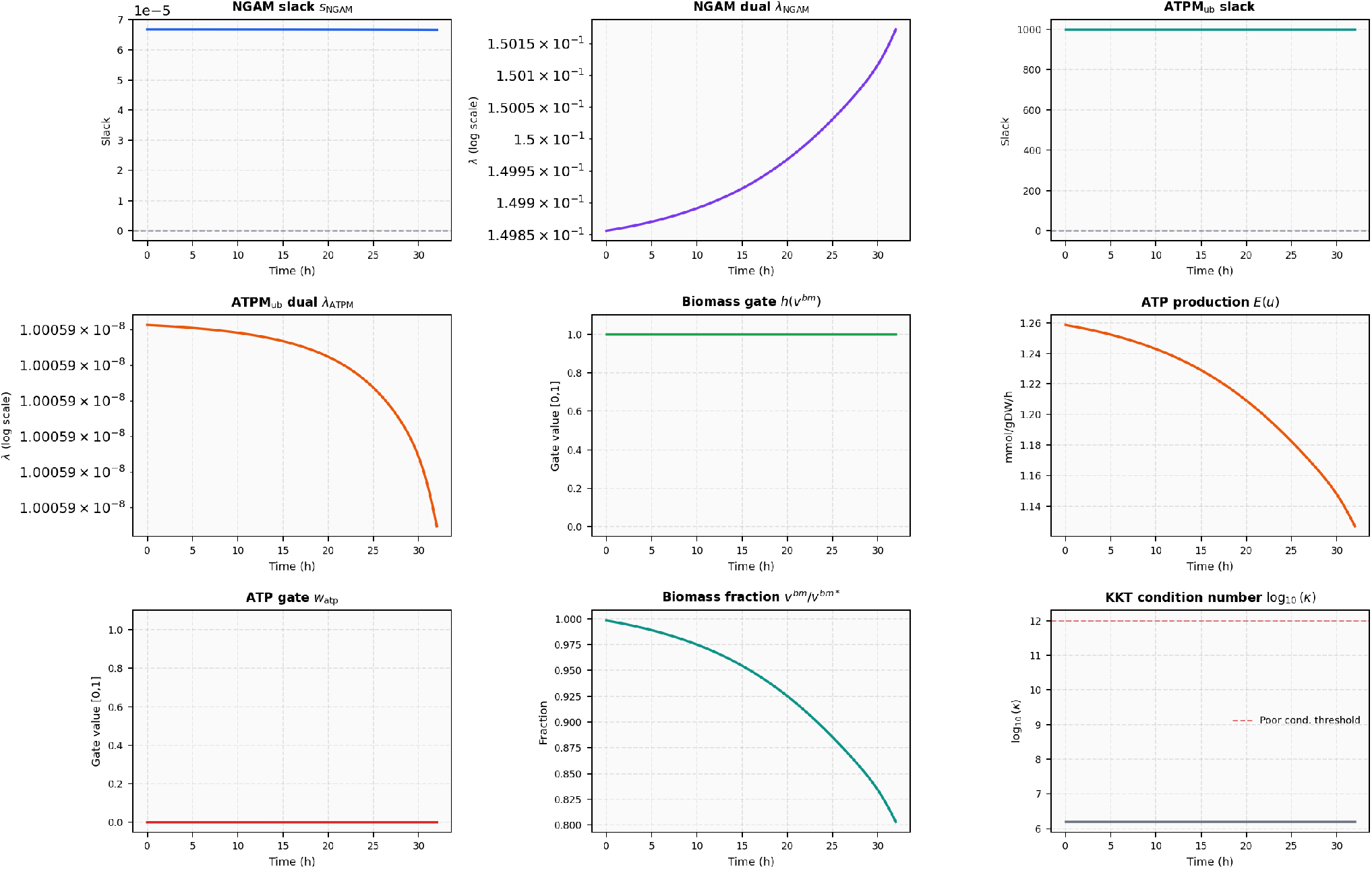
Diagnostics for the successful simulation (exp. 1974, seed 0). All slacks remain strictly positive, gating functions stay inactive, and the KKT condition number stays well below 10^12^ throughout the 32 h batch.

The NGAM lower-bound slack *s*_NGAM_ (top left) remains at ~10^−5^, reflecting the fixed barrier parameter *µ* = ~10^−5^: the constraint is nearly active throughout but never violated. Its dual *λ*_NGAM_ (top centre) grows gradually, indicating that the ATP maintenance requirement becomes relatively more binding as substrates are consumed and metabolic flexibility narrows. The NGAM upper-bound slack (top right) stays large (≫1), confirming that the upper-bound constraint is not active.

The biomass gate *h*(*v*^*bm*^) (centre left) remains at 1 throughout the full 32 h batch: biomass flux never drops far enough to activate the gate, which is the expected behaviour in a healthy simulation. The ATP production flux *E*(*u*) (centre) declines gradually as carbon sources are depleted. The ATP maintenance gate *w*_atp_ (bottom left) stays at zero, confirming that ATP feasibility is never threatened. The biomass fraction *v*^*bm*^*/v*^*bm*\*^ (bottom centre) declines monotonically from near 1 as the substrate pool is consumed, tracking the shrinking feasible polytope.

The KKT condition number log_10_(*κ*) (bottom right) remains well below the 10^12^ threshold throughout, confirming that the linear system (7) is numerically well-conditioned for this prior draw.

## Appendix E FAILED SIMULATION EXAMPLE

Figures E.1 and E.2 show the same experiment as the successful case (exp. 1974) but with a different prior draw (seed 3), which crashes at *t* ≈ 19.4 h. Using the same experiment isolates the failure to the parameter draw rather than the medium composition, making the contrast interpretable.

**Fig. E.1.**
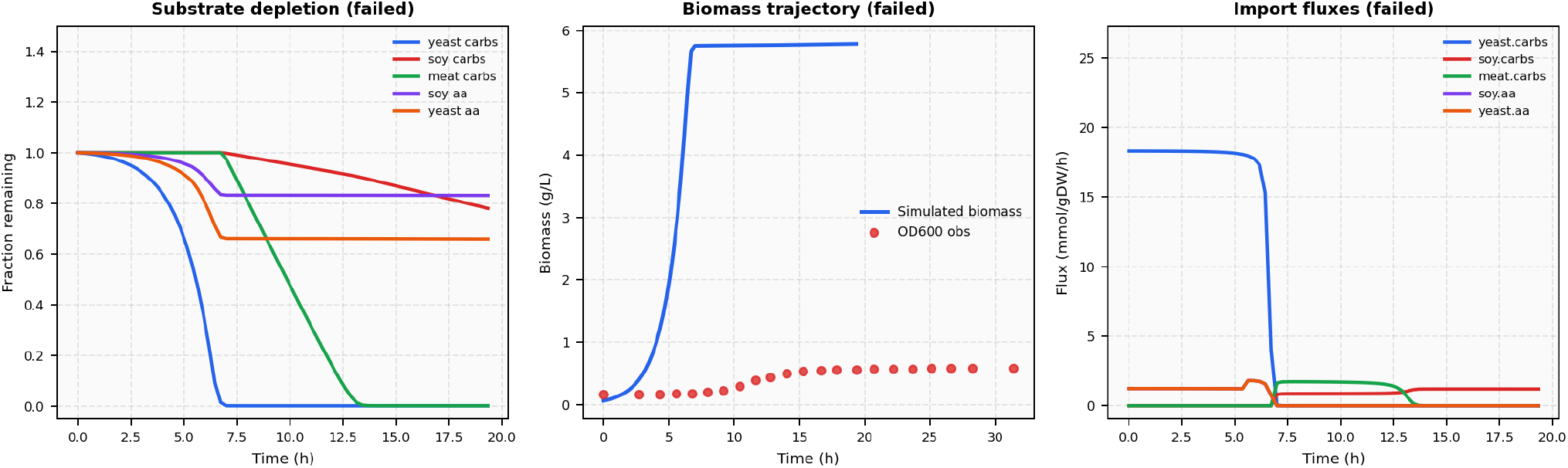
State trajectories for a failing simulation (exp. 1974, seed 3; final time *t* = 19.4 h). The same experiment as Figure 7 with a different prior draw, isolating the failure to the parameter sample. Biomass grows normally to a peak at *t* ≈ 7 h then collapses; import fluxes drop sharply at the same time as the polytope becomes infeasible under this parameter draw.

The state trajectories (Figure E.1) show biomass growing normally in a sigmoid shape up to *t* ≈ 7 h, where it peaks and then falls back toward zero. Import fluxes collapse sharply at the same time. The OD600 observations (red dots) are identical to Figure 7; the simulated trajectory diverges because this parameter draw implies different kinetics.

The diagnostic panel (Figure E.2) reveals the failure mechanism. The NGAM slack *s*_NGAM_ (top-left) is *negative from t* = 0: the parameter draw demands more ATP maintenance than the initial medium can supply, so the softmin gate is engaged from the outset. Despite this, the simulation proceeds as long as ATP production *E*(*u*) (centre-right) remains non-zero. At *t* ≈ 7 h, substrate depletion drives *E*(*u*) to collapse abruptly to zero; the ATPM upper-bound slack (top-right) simultaneously falls to zero and the NGAM dual (top-centre) jumps by several orders of magnitude, indicating the constraint has become strongly active. The biomass gate *h*(*v*^*bm*^) (centre-left) and biomass fraction (bottom-centre) both drop sharply to zero at *t* ≈ 7.5 h as biomass flux becomes unattainable. The ATP maintenance gate *w*_atp_ (bottom-left) rises toward 1 around *t* ≈ 7 h in response to the emerging infeasibility, but is unable to restore feasibility once ATP production has fully collapsed. Following this the KKT system becomes progressively ill-conditioned; the condition number (bottom-right) crosses the 10^12^ threshold around *t* ≈ 15 h and continues rising until the integrator exhausts its step budget at *t* = 19.4 h. The failure cascade is: infeasible NGAM from draw ⇒ ATP maintenance gate activates ATP production collapses ⇒ polytope contracts ⇒ ill-conditioned KKT system ⇒ integrator failure.

**Fig. E.2.**
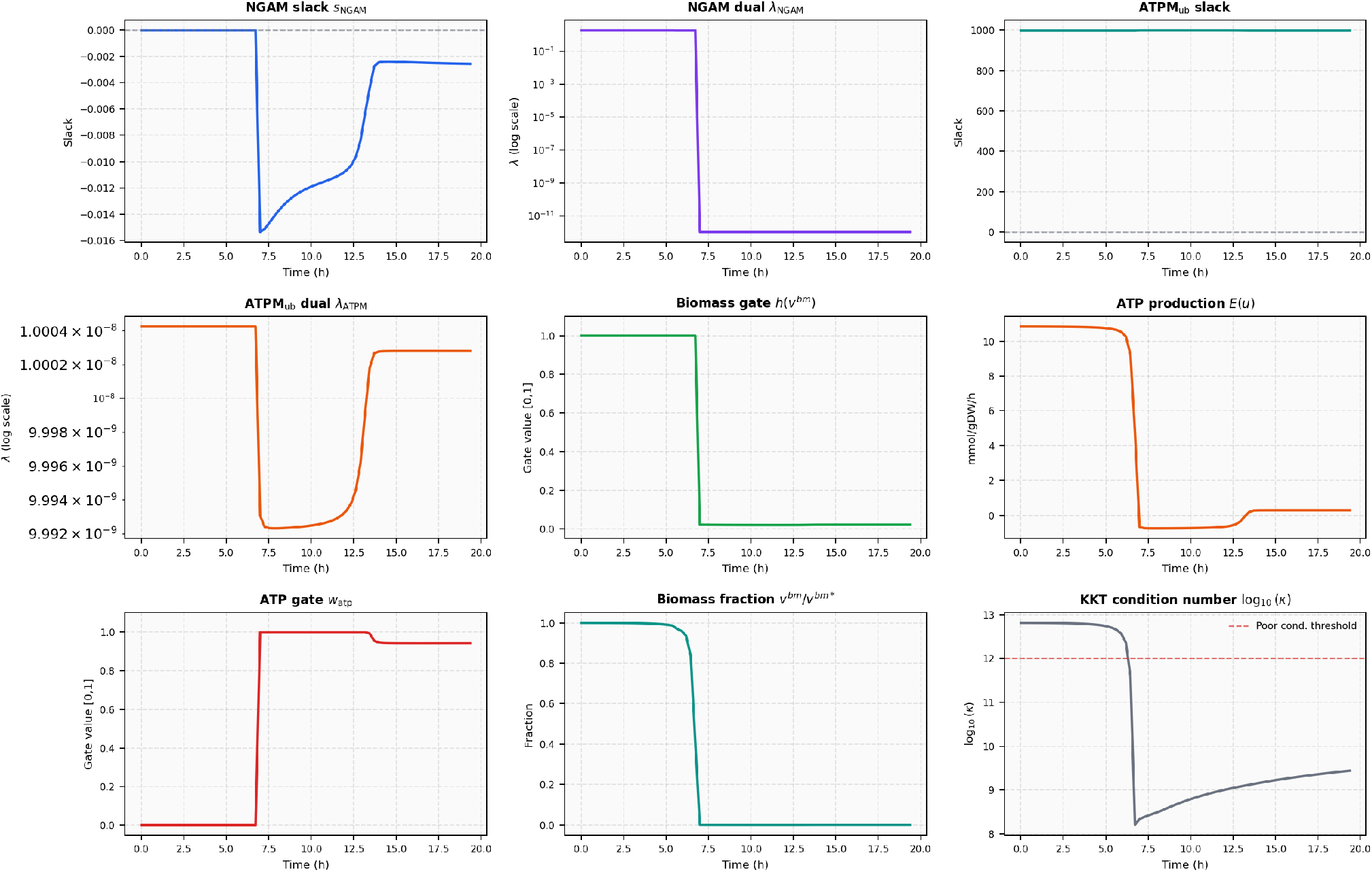
Diagnostics for the failing simulation (exp. 1974, seed 3). NGAM slack negative from *t* = 0; ATP maintenance gate activates at *t* ≈ 7 h then fails to restore feasibility; biomass gate and fraction collapse at *t* ≈ 7.5 h; KKT condition number exceeds 10^12^ near *t* = 15 h and integrator terminates at *t* = 19.4 h. fraction of prior draws imply parameter combinations that stress the KKT system.

## Appendix F COMPLETION RATE SWEEPS

Two sweeps characterise the completion rate of the simulator across parameter and experiment variation.

**Sweep A** (Figure F.1) fixes exp. 1974 and varies the prior draw seed from 0 to 29. 25 of 30 draws (83%) complete the full 32 h batch. The 5 failures occur at seeds 3, 8, 12, 21, and 23. All fail due to ODE step-size collapse: the NGAM slack approaches zero under those parameter combinations, the barrier Hessian diverges, and the integrator exhausts its step budget. The failing seed 3 is examined in detail in Appendix E. The spread of failure seeds across the range suggests that failures are not concentrated in a corner of parameter space but arise from a minority of draws whose kinetic constants imply marginal ATP feasibility for this particular medium.

**Fig. F.1.**
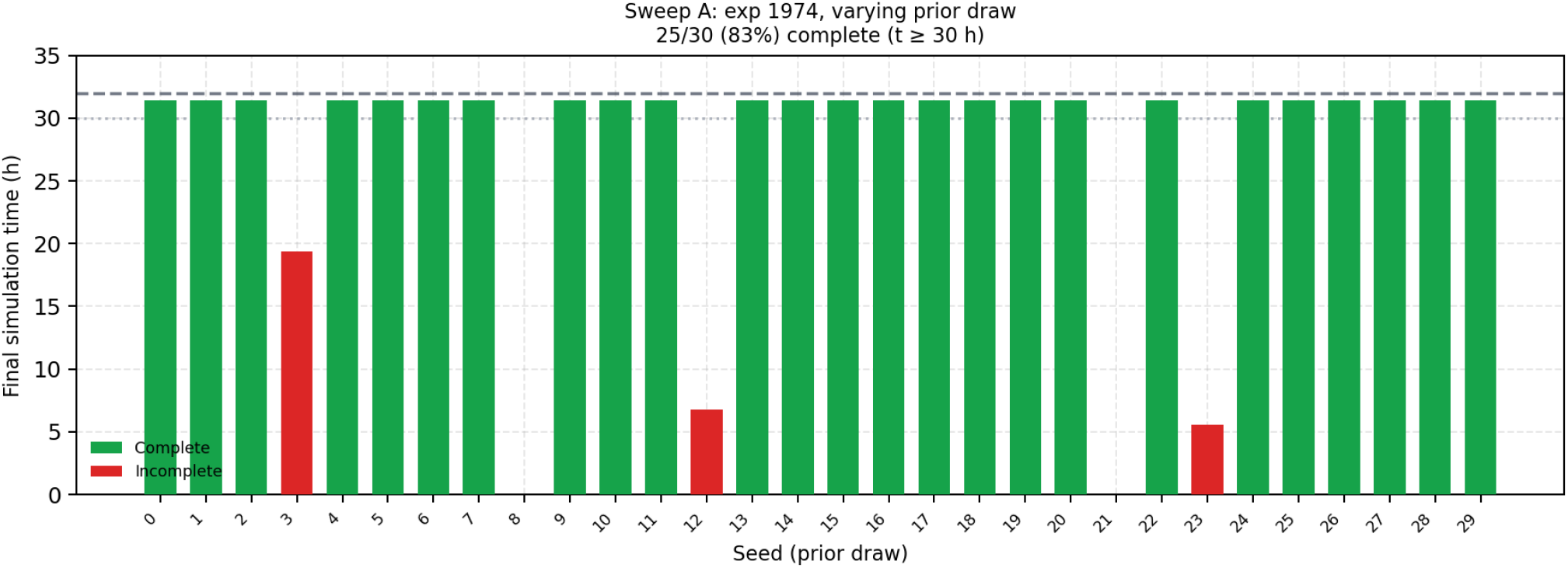
Sweep A: completion status for 30 prior draws at fixed experiment 1974. Green bars reach *t* ≥ 30 h; red bars terminate early with final time labelled. 25/30 (83%) complete.

**Sweep B** (Figure F.2) fixes seed 0 and samples 30 evenly-spaced experiment indices across the full dataset of ~2 100 fermentations. 29 of 30 experiments (97%) complete. The single failure (exp. 793) terminates at *t* = 4.3 h. The high completion rate across experiments with a fixed parameter draw indicates that the seed-0 parameters are relatively robust, and that the primary source of completion-rate variation is the prior rather than the medium composition. The moderate failure rate in Sweep A (17%) compared to Sweep B (3%) supports this interpretation: most medium compositions are numerically tractable, while a non-trivial

**Fig. F.2.**
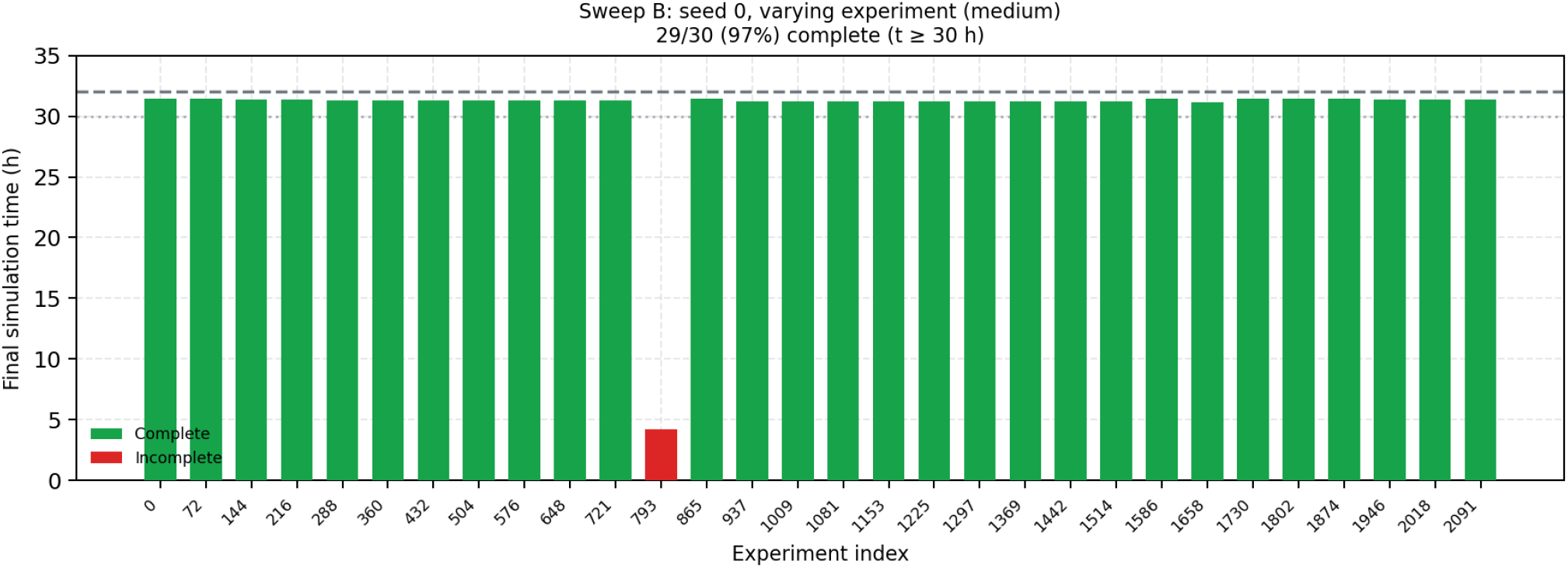
Sweep B: completion status for 30 evenly-spaced experiments with fixed prior draw (seed 0). 29/30 (97%) complete; the single failure (exp. 793) terminates at *t* = 4.3 h.

Together the sweeps suggest that targeted prior regularisation— penalising draws that push the NGAM slack towards zero at batch end—may be a more efficient route to improving completion rate than further engineering of the ODE solver.

## Appendix G END-TO-END GRADIENT VALIDATION

We tested gradient validity for all 54 (experiment, seed) pairs that completed successfully in Sweeps A and B (Section F). For each pair, we ran `simulate_ode` in non-debug mode and computed ∇_*θ*_*L* via `jax.value_and_grad`, where *L* = ∑_*t*_(*c*^*bm*^(*t*; *θ*))^2^ is a scalar loss over the biomass trajectory.

Figure G.1 shows the results: all 54 pairs fail. The error is consistent across pairs—an `EquinoxRuntimeError` from the linear solver returning non-finite output during the adjoint backward pass. The root cause is the KKT matrix *M* in (7):

**Fig. G.1.**
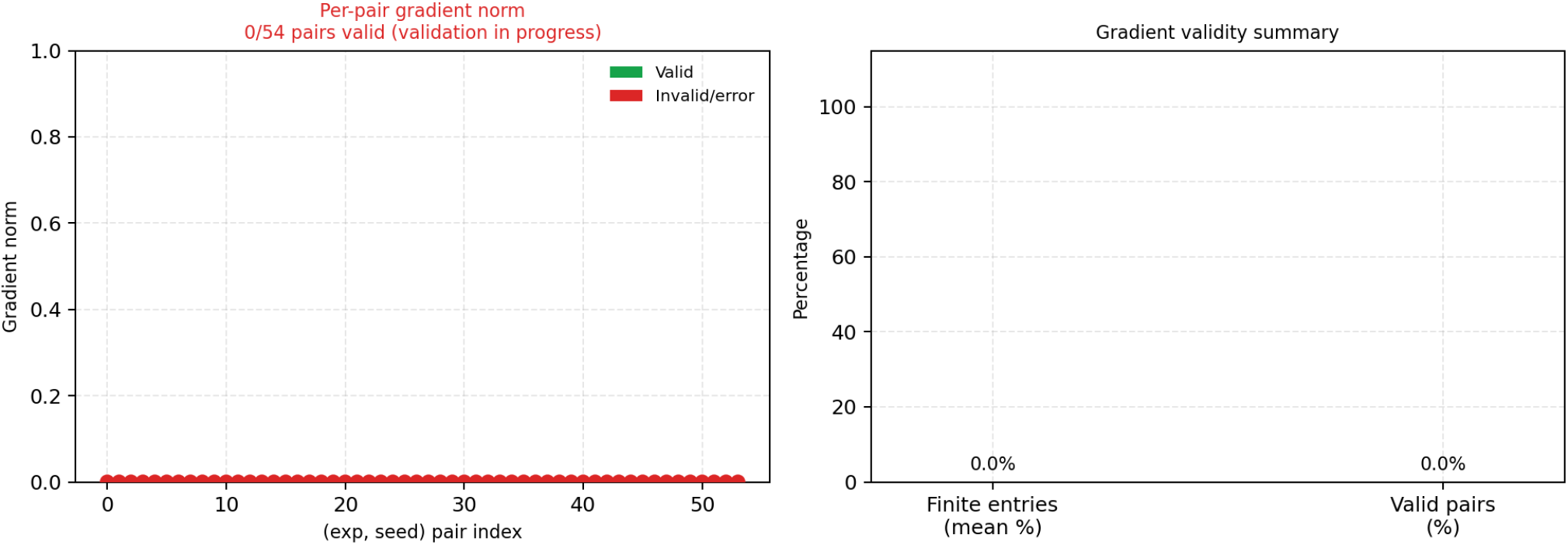
End-to-end gradient validity across all 54 (experiment, seed) pairs that completed successfully in both sweeps. All 54 pairs fail: the adjoint backward pass through the KKT linear system returns non-finite values due to ill-conditioning of *M* (G.1) as slack variables approach zero during backward integration.

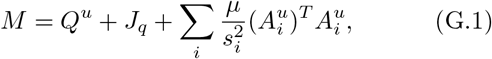

whose condition number *κ*(*M*) grows without bound as slack variables *s*_*i*_ approach zero during backward integration. While the forward pass can tolerate mild ill-conditioning (the step-size controller rejects steps where the solver fails), the adjoint backward pass has no such safeguard and returns NaN as soon as the linear solve fails.

Two fixes are known and under implementation:

1. **KKT regularisation during the adjoint pass**: replace *M* with *M* +*ϵI* for a small diagonal regulariser *ϵ >* 0 that is applied only during the backward pass, leaving the forward dynamics unchanged.
2. `stop_gradient` **on dual variables**: the dual variables *λ* are algebraically determined by the primal state and do not need to be differentiated; applying `stop_gradient` to *λ* before it enters the backward KKT solve removes the ill-conditioned path entirely.

Either fix is expected to restore finite gradients for the majority of pairs. Once implemented, gradient validity will be re-evaluated on the full sweep set before variational inference training begins.

